# Cost-effective long-read assembly of a hybrid *Formica aquilonia* × *Formica polyctena* wood ant genome from a single haploid individual

**DOI:** 10.1101/2021.03.09.434597

**Authors:** Pierre Nouhaud, Jack Beresford, Jonna Kulmuni

## Abstract

*Formica* red wood ants are a keystone species of boreal forest ecosystems and an emerging model system in the study of speciation and hybridization. Here we performed a standard DNA extraction from a single, field-collected *Formica aquilonia* × *Formica polyctena* haploid male and assembled its genome using ∼60× of PacBio long reads. After polishing and contaminant removal, the final assembly was 272 Mb (4,687 contigs, N50 = 1.16 Mb). Our reference genome contains 98.5% of the core Hymenoptera BUSCOs and was scaffolded using the pseudo-chromosomal assembly of a related species, *F. selysi* (28 scaffolds, N50 = 8.49 Mb). Around one third of the genome consists of repeats, and 17,426 gene models were annotated using both protein and RNAseq data (97.4% BUSCO completeness). This resource is of comparable quality to the few other single individual insect genomes assembled to date and paves the way to genomic studies of admixture in natural populations and comparative genomic approaches in *Formica* wood ants.

## INTRODUCTION

Despite their small size, red wood ants (subgenus *Formica sensu stricto*, or *F. rufa* group, hereafter wood ants) are heavyweights of boreal ecosystems. These social insects build massive interconnected nest mounds forming supercolonies of several million individuals, covering up to 2 km^2^ (Stockan, Robinson, Trager, Yao, & Seifert, 2016). Wood ants are considered keystone species which play a role in nutrient cycling (Frouz, Jílková, & Sorvari, 2016), predator–prey dynamics or plant growth (Robinson, Stockan, & Iason, 2016), to name a few.

Wood ant genomics have so far mostly focused on supercoloniality, which is an extreme form of sociality. While the canonical ant colony is headed by a single queen (monogyny) and occupies a unique nest, in wood ant supercolonies a single nest may contain dozens to hundreds of unrelated egg-laying queens (polygyny, Pamilo, 1993). This social polymorphism is governed by a supergene maintained across species which diverged 40 Mya (Brelsford et al., 2020; Purcell, Brelsford, Wurm, Perrin, & Chapuisat, 2014).

Wood ants have undergone recent radiation (Goropashnaya, Fedorov, Seifert, & Pamilo, 2012) and represent a promising system for the study of speciation and hybridization. This process is ubiquitous across living organisms and haplodiploids (organisms for which one sex is haploid and the other, diploid) such as wood ants can answer some key questions in admixture research which are difficult to study in standard diploid organisms (Nouhaud, Blanckaert, Bank, & Kulmuni, 2020). The best-characterised case is the occurrence of natural hybrids between *F. aquilonia* and *F. polyctena* in Southern Finland. Two hybrid lineages coexist in a single population (Kulmuni, Seifert, & Pamilo, 2010), where introgression between lineages is sex-specific but could be modulated by external factors (Kulmuni et al., 2020).

Currently, no high-quality reference genome is available for any species of the *Formica s. str*. subgenus. Kulmuni et al. (2020) assembled a draft genome using poolseq data from a hybrid *F. aquilonia* × *F. polyctena* population, but the assembly is highly fragmented (> 300k contigs, N50 < 2 kbp). At a broader phylogenetic scale, among palaearctic *Formica* species, two genomes are available for the *Coptoformica* and *Serviformica* subgenera, respectively *F. exsecta* (Dhaygude, Nair, Johansson, Wurm, & Sundström, 2019) and *F. selysi* (Brelsford et al., 2020). However, divergence is around 10% between each subgenus pair (Goropashnaya et al., 2012), which precludes using either of these genomes as references when studying members of the *Formica s. str*. clade.

While PacBio DNA input requirements have for a long time hindered the individual sequencing of small organisms, a modified SMRTbell library construction protocol was recently used to build a reference genome from a single *Anopheles* mosquito (Kingan, Heaton, et al., 2019). Few other recent examples demonstrate that high-quality arthropod genomes can now be built from a single individual (fruit fly: Adams et al., 2020; lanternfly: Kingan, Urban, et al., 2019; braconid wasp: Ye et al., 2020). Here, we assemble the genome of a single haploid, hybrid *F. aquilonia* × *F. polyctena* male using PacBio sequencing. As sexuals from these species are relatively big (∼20 mg), we could apply a cost-effective, standard extraction protocol to obtain high-molecular weight DNA from a single individual. The contigs were anchored against the *F. selysi* chromosomal assembly after contamination removal, and the genome was annotated using both RNAseq and protein data. Overall, the quality of the assembly and its annotation are on par with other single individual insect genomes published to date as well as other sequenced ant genomes.

## MATERIALS & METHODS

### Sampling

All individuals used in the present study were sampled from the Långholmen population in Southern Finland (59°50’59.9”N, 23°15’03.3”E) in Spring 2018. This population has been characterized as a hybrid between *F. aquilonia* and *F. polyctena* using both genetic markers and morphological data (Kulmuni et al., 2010; Seifert, Kulmuni, & Pamilo, 2010). The Långholmen population is a supercolony consisting of two genetic lineages of hybrid origin (R and W; (Kulmuni & Pamilo, 2014; Kulmuni et al., 2010), which show moderate genetic differentiation (*F*_ST_ ≈ 0.105, Kulmuni et al., 2020).

For long-read sequencing, a single haploid male was collected from the FAu2014a nest (W lineage) in Spring 2018. Two males and two unmated gynes (queens) from the same nest and lineage were also sampled at the same time for polishing purposes (short-read sequencing, see below). All samples were collected in individual sterile tubes and flash-frozen on the field. For RNA sequencing, sexual larvae were collected from multiple R and W nests in the same population in May 2014, measured and put in individual tubes before flash-freezing in the laboratory within 24 hours of collection (Beresford et al., unpublished). All samples were stored at -80ºC without any buffer.

### Long-read sequencing

For both PacBio and Illumina DNA sequencing, all steps were carried by Novogene (Hong Kong) as part of the Global Ant Genomics Alliance (GAGA, Boomsma et al., 2017). DNA from a single haploid male was extracted using a Sodium Dodecyl Sulfate (SDS) protocol, and a SMRTbell library was prepared using the SMRT bell Template Prep Kit 1.0-SPv3 (Pacbio, 100-991-900). DNA quantification was performed using a Qubit fluorometer (Thermo Fisher) and purity was assessed with an agarose gel electrophoresis. The extraction from a single male yielded 9.89 μg of DNA, at a concentration of 86 ng.μl^-1^ (A_260/280_ = 1.76, A_260/230_ = 1.20). DNA fragmentation was assessed through an Advanced Analytical Fragment Analyzer (AATI, mean size: 18317 bp) prior to size selection (BluePippin, Sage Sciences, cutoff: 10 kb). The sample was loaded onto four SMRT cells with the Sequel Sequencing Kit 2.0 following PacBio recommendations and sequenced on a PacBio Sequel platform.

### Short-read DNA sequencing

Since accuracy of long-read data is lower than short-read data (e.g., Koren et al., 2012; but see Wenger et al., 2019), Illumina data was generated to correct spurious base calls. For the four samples used for these polishing purposes, DNA was extracted from whole bodies with a SDS protocol and libraries were constructed using NEBNext DNA Library Prep Kits (New England Biolabs). Whole-genome sequencing was performed on Illumina Novaseq 6000 (paired-end mode, 150 bp), after which raw Illumina reads and adapter sequences were trimmed using Trimmomatic (v0.38; parameters LEADING:20 TRAILING:20 MINLEN:50; Bolger, Lohse, & Usadel, 2014).

### Whole-genome assembly

We assessed the performance of two long-read assemblers, Canu (v1.8, Koren et al., 2017) and wtdbg2 (v2.5, Ruan & Li, 2020). We assumed a haploid genome size of 323 Mb, which is the mean size estimated from five species of the Formicinae subfamily by flow cytometry (Tsutsui, Suarez, Spagna, & Johnston, 2008). Canu was run with default parameters, except that the maximum allowed difference threshold was adapted to Sequel data (correctedErrorRate=0.085), following Canu’s FAQ. For wtdbg2, a first run was performed using settings optimised for Sequel data and genome sizes below 1 Gb (preset 2: -p 0 -k 15 -AS 2 -s 0.05) but selecting all subread lengths (-L 0). Based on the subread distribution, a second run was performed with the same preset, but selecting only subreads above 10 kb (-L 10000). For each assembly we assessed completeness using BUSCO (v4.0.5, Seppey, Manni, & Zdobnov, 2019) with the Hymenoptera ODB gene set v10.

The canu assembly contained 338 Mb in 3,633 contigs (assuming a haploid genome size of 323 Mb, NG50 = 283 kb), the wtdbg2 assembly using all subreads totaled 349 Mb in 11,615 contigs (NG50 = 71 kb) and running wtdbg2 only with subreads greater than 10 kb (∼44×) yielded a 280 Mb assembly with 5,098 contigs (NG50 = 689 kb). Because of a large fraction of missing BUSCOs, the wtdbg2-all assembly was discarded (supplementary table 1). The completeness of both canu and wtdbg2-10k assemblies were comparable (97.5% vs. 97.1%, respectively) despite stark differences in total sizes (338 Mb vs. 280 Mb, respectively). However, the BUSCO duplication rate was much higher for the canu assembly (5.7% vs. 0.5%). This suggests that the canu assembly may contain duplicated regions, which could in turn inflate its size. Interestingly, while the average genome size for the Formicinae subfamily was estimated at 323 Mb by flow cytometry (Tsutsui et al., 2008), recent genome projects within the *Formica* genus documented genome sizes much closer to our 280 Mb estimate, with 278 Mb for *F. exsecta (Dhaygude et al*., *2019)* and 290 Mb for *F. selysi* (Brelsford et al., 2020). Based on this observation, plus the assembly statistics and BUSCO score (supplementary table 1), we concluded that the wtdbg2-10k assembly was the best model. The next steps were only performed on this assembly.

### Assembly polishing

To avoid incorporating sequencing errors in our final assembly (Watson & Warr, 2019), we polished our contigs using Racon (v1.4.10, Vaser, Sović, Nagarajan, & Šikić, 2017). We ran four polishing iterations with the PacBio data, followed by two iterations with the Illumina resequencing data (all four individuals pooled), always keeping unpolished sequences in the output (parameter -u). For each iteration, alignment was performed using minimap2 (v2.17, Li, 2016, using parameters -x map-pb for PacBio and ax sr for Illumina data, respectively).

### Contaminant removal & mitochondrial genome identification

The assembly was assessed for contaminants with BlobTools (v1.1.1, Laetsch & Blaxter, 2017). Coverage files were obtained using minimap2 for both Canu-corrected PacBio subreads and the four resequenced individuals. Taxonomic partitioning of contigs was carried through BLAST against the

NCBI non-redundant database. The contig containing the mitochondrial genome was identified based on BlobTools results (lower GC proportion compared to the rest of the genome and high sequencing depth, supplementary fig. 1) and was further validated by BLAST of the *F. selysi* mtDNA sequence (Brelsford et al., 2020) against the whole assembly. *Formica* ants carry *Wolbachia* endosymbionts (Viljakainen, Reuter, & Pamilo, 2008) and horizontal gene transfer (HGT) has been previously characterized in *F. exsecta (Dhaygude et al*., *2019)*. To avoid classifying ant contigs impacted by HGT as contigs of endosymbiont origin, we blasted the closest *Wolbachia* genome (NCBI accession PRJNA436771) against our assembly and manually inspected these results in conjunction with coverage profiles and the physical location of Hymenoptera BUSCO hits (v4.0.5, Seppey et al., 2019).

### Anchoring of contigs to pseudo-chromosomes

Our polished, ant nuclear contigs were coalesced into pseudo-chromosomes with RaGOO (v1.1, Alonge et al., 2019), using the *F. selysi* reference genome (Brelsford et al., 2020) as a guide. To evaluate RaGOO’s performance, we also aligned contigs against *F. selysi* pseudo-chromosomes using the nucmer aligner from MUMmer (v4.0.0beta2, Marçais et al., 2018). Delta files from nucmer were processed using the DotPrep.py script (https://dnanexus.github.io/dot/DotPrep.py, last accessed 20.10.2020) and alignments were visualised using Dot (https://dot.sandbox.bio/, last accessed 20.10.2020). A large portion (6 Mb) of Scaffold 10 in *F. selysi* mostly contained highly repetitive alignments (see also Figure 1 from Brelsford et al., 2020). This region was removed from the *F. selysi* assembly before a second RaGOO run was performed. The gap size was set to 100 (100×N). All remaining, unanchored contigs were scaffolded as a single Scaffold 0. Of note, both parental *species F. aquilonia* and *F. polyctena* have 26 chromosomes (n = 26, Rosengren & Rosengren, 1980), while *F. selysi* has 27 (*n* = 27). Unfortunately, the breakpoint could not be identified on the sole basis of our long-read data. Our assembly contains 27 pseudo-chromosomes instead of the 26, which is the correct karyotype for both parental species.

**Fig. 1.**
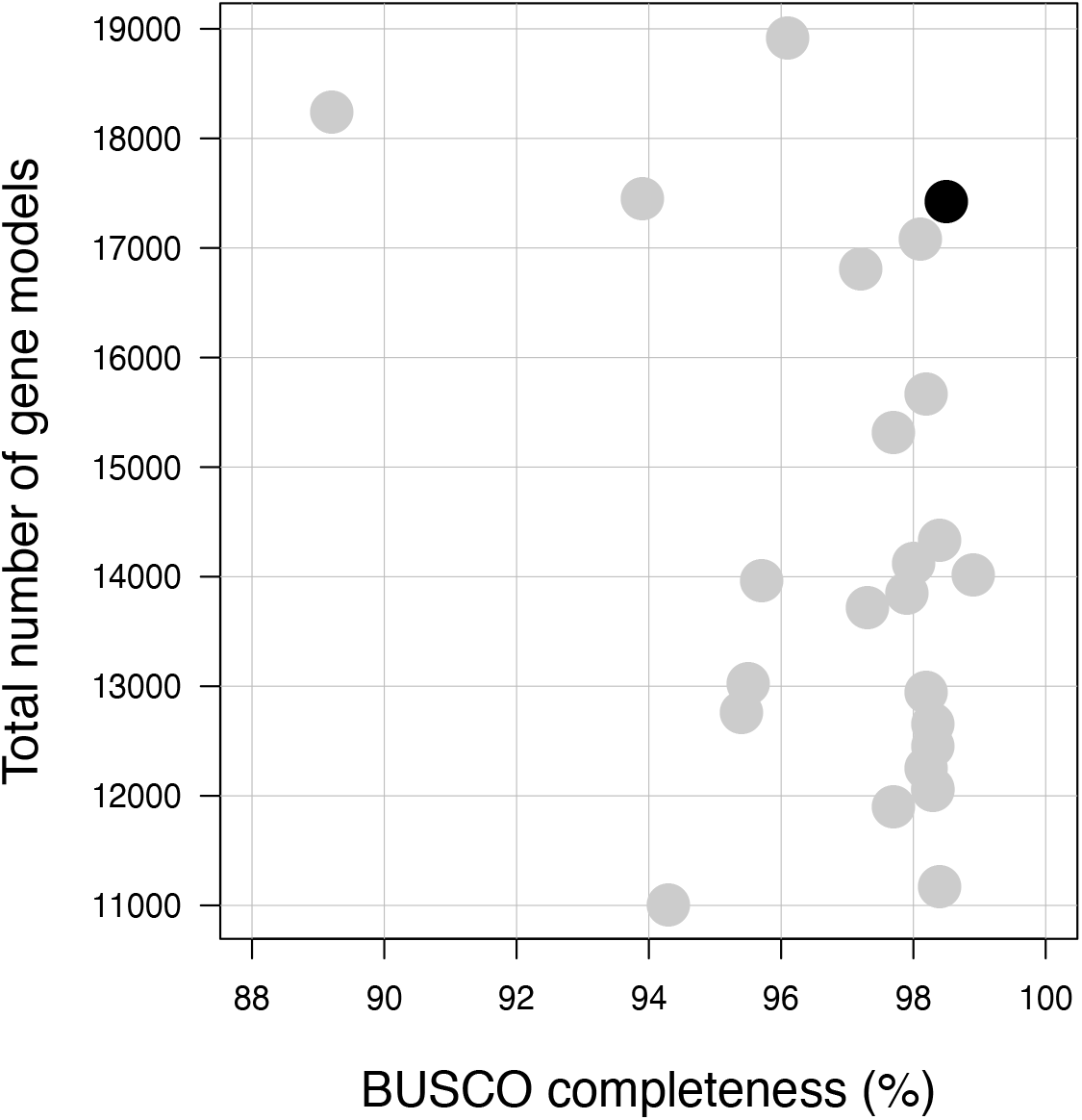
Total number of gene models as a function of BUSCO genome completeness metrics in ant genomes for which annotations are available on NCBI (*n* = 24, light gray) and the assembly of this study (black). Detailed statistics are shown in supplementary table 3.

### Annotation of repeat sequences

Transposable elements (TEs) were annotated using the Dfam TE Tools Container (v1.1, https://github.com/Dfam-consortium/TETools, last accessed 20.10.2020). A de novo consensus library was built with Repeatmodeler 2 (Flynn et al., 2020) and used to mask TE sequences in our assembly using Repeatmasker (Smit, Hubley, & Green, 2013).

### RNA sequencing

For annotation purposes, RNAseq data was generated for nine individuals originating from six nests in the Långholmen population (R: nest FA4, 3 individuals and W: nest FA15, 1 ind.; FA17, 2 inds.; FA25, 1 ind.; FA35, 1 ind.; FAu2014a, 1 ind.). These individuals were at different larval stages and total RNA was extracted from whole bodies using an ALLPrep DNA/RNA Mini Kit (Qiagen) following manufacturer’s instructions. Individual RNA qualities were assessed using a Bioanalyzer (Agilent 2100). Libraries were constructed using NEBNext Ultra RNA Library Prep Kits and samples were sequenced on an Illumina NextSeq platform (paired-end mode, 150 bp) at the Biomedicum Functional Genomics Unit (FuGU, University of Helsinki). Raw reads were trimmed using Trimmomatic (v0.38, parameters LEADING:20 TRAILING:20 SLIDINGWINDOW:5:20 MINLEN:50 Bolger et al., 2014) and unpaired reads were discarded. Approximately 5.60 million 150bp paired reads were randomly sampled per individual and combined into two (Forward and Reverse) FASTQ files, totaling 50 million paired reads over all individuals.

### Genome annotation

We annotated the genome with the Braker2 pipeline (Brůna, Hoff, Lomsadze, Stanke, & Borodovsky, 2020; v2.1.5, Hoff, Lomsadze, Borodovsky, & Stanke, 2019). Both RNAseq- and protein-derived hints were used to train GeneMark-ETP, which predictions were in turn used to train Augustus and obtain the final gene set. All protein data available for Arthropoda were downloaded from OrthoDB (v10, Kriventseva et al. 2019, https://v100.orthodb.org/download/odb10_arthropoda_fasta.tar.gz, last accessed 22.07.2020) and aligned using ProtHint. This dataset contains 2.6 million sequences and encompasses 170 species, including 40 of the same order (Hymenoptera) and 17 of the same family (Formicidae). RNAseq data produced above was aligned against the hard-masked genome using STAR (v2.7.2, Dobin et al., 2013), and secondary alignments were removed with SAMtools (v1.10, Li et al., 2009). After the Braker2 run, protein sequences of all gene models not supported by at least one hint were blasted against the Uniprot database (UniProt Consortium, 2019) and all models without any hit on Aculeata (wasps, bees and ants) were discarded from the final gene set. Finally, functional annotation was carried with EnTAP (v0.10.3, Hart et al., 2020) using the EggNOG (Huerta-Cepas et al., 2016), Uniprot and RefSeq databases.

## RESULTS AND DISCUSSION

### Genome sequencing & assembly

We generated 2,547,044 subreads on the PacBio Sequel, suming to 21.8 Gb of data (∼68×). Half of the subreads were longer than 11.5 kb (NR50), with a mean length of 8.55 kb.

Running wtdbg2 only with subreads greater than 10 kb (∼44×) yielded a 280 Mb assembly with 5,098 contigs (assuming a haploid genome size of 323 Mb, NG50 = 689 kb, supplementary table 1). Interestingly, while the average genome size for the Formicinae subfamily was estimated at 323 Mb by flow cytometry (Tsutsui et al., 2008), recent genome projects within the *Formica* genus documented genome sizes much closer to our 280 Mb estimate, with 278 Mb for *F. exsecta (Dhaygude et al*., *2019)* and 290 Mb for *F. selysi (Brelsford et al*., *2020)*. Based on this observation (similar assembly sizes for different *Formica* species) and BUSCO metrics, we concluded that our assembly had a sufficiently high level of completeness.

After polishing using both long (four iterations) and short reads (two iterations), the BUSCO score reached 98.5% for complete single-copy orthologs (table 1) while the total size of the assembly reduced to 276 Mb.

**Table 1.** Assembly & annotation metrics. *: scaffold statistics computed after excluding both the mitochondrial genome and Scaffold 0, which contains all unanchored contigs (59 Mb, “total unanchored length”).

|  |  |
| --- | --- |
| **Genome assembly** |  |
| BUSCO v4.0.5 genome score | C:98.5%[S:97.9%,D:0.6%], F:0.4%,M:1.1%,n:5991 |
| Number of contigs | 4,687 |
| Contig N50 (bp) | 1,163,114 |
| Shortest contig (bp) | 117 |
| Longest contig (bp) | 4,650,116 |
| Average contig length (bp) | 58,036 |
| Total contig length (bp) | 272,015,305 |
| Number of scaffolds | 28 |
| Scaffold N50* (bp) | 8,490,488 |
| Shortest scaffold (bp) | 3,646,393 |
| Longest scaffold* (bp) | 14,915,360 |
| Average scaffold length* (bp) | 7,887,222 |
| Total scaffold length (bp) | 272,497,664 |
| Total unanchored length (bp, fraction) | 59,526,201 (21.8%) |
| GC content | 36.3% |
| N fraction | 0.17% |
| **Genome annotation** |  |
| BUSCO v4.0.5 protein score | C:97.4%[S:96.8%,D:0.6%], F:1.4%,M:1.2%,n:5991 |
| Total number of gene models | 17,426 |
| Mean gene length (bp) | 5,524 |
| Average number of exons per gene | 5.80 |
| Number of models with RNAseq support (fraction) | 11,956 (68.6%) |
| Number of isoforms | 19,226 |
| Average number of isoforms per gene | 1.10 |
| Cumulative gene length (bp, fraction) | 78,835,002 (29.0%) |
| Cumulative exon length (bp, fraction) | 27,442,032 (10.1%) |
| **Repeat annotation** |  |
| Fraction of genome masked | 32.01% |
| Interspersed repeats, total fraction | 28.44% |
| Retroelements (class I) | 6.39% |
| LINEs | 1.47% |
| Gypsy/DIRS1 | 2.72% |
| DNA transposons (class II) | 3.56% |
| Unclassified | 18.50% |
| Simple repeats | 2.59% |

Almost 92% (4,688) of the 5,098 contigs were assigned to Arthropoda, while 82 contigs were assigned to Proteobacteria (supplementary fig. 1). *Formica* ants harbor *Wolbachia* endosymbionts (Viljakainen et al., 2008), and HGT between *Wolbachia* and the ant nuclear genome has been characterized (Dhaygude et al., 2019). Through manual curation we assigned 76 contigs to *Wolbachia* (total size = 1,786,664 bp, N50 = 33.4 kb) and six contigs of the nuclear ant genome as putative HGTs. Overall, the contamination removal step decreased the nuclear ant genome size to 272,015,305 bp.

Finally, we anchored 78.2% (213 Mb, table 1) of our assembly to the 27 pseudo-chromosomes of the *F. selysi* genome, a fraction similar to that of the original *F. selysi* study (78.3% of the assembly assigned to pseudo-chromosomes, see Table S3 in Brelsford et al., 2020).

### Genome annotation

Overall, 32% of the sequence was masked with Repeatmasker, most of the repeats being unclassified (18.5%), 6.39% being retroelements and 3.56% being DNA transposons (table 1). The vast majority of repeats were located on unanchored contigs (supplementary fig. 2).

The initial gene set contained 30,068 gene models, which is far superior to what has been documented in ants (∼17,000 gene models, Gadau et al., 2012). Among these models, 14,287 (47.5%) were not supported by any protein or RNAseq hint. Moreover, the size of these hint-less models was much shorter than hint-supported models. As we suspected an overprediction problem (which was also observed for alternative Braker2 runs, supplementary table 2), we only kept hint-less models if their protein sequences had a blast hit against Aculeata in Uniprot, which reduced the total set from 30,068 to 17,426 gene models (15,781 with hints plus 1,645 recovered after blast). Overall, 19,226 mRNAs were identified, among which 15,664 (81.5%) were functionally annotated with EnTAP. From these, 63.4% of the proteins had their best hit within ant species or *Drosophila melanogaster* (supplementary fig. 3). The completeness of this final gene set assessed with BUSCO was good (protein mode: 97.4%, table 1) and our assembly showed a level of completeness comparable to other ant genomes annotated so far (fig. 1, supplementary table 3).

## Conclusions

Here we report a pseudo-chromosome-level genome assembly for a single hybrid *F. aquilonia* × *F. polyctena* haploid male using a simple and cost-effective extraction protocol. The final assembly sums to 272 Mb, of which 78.2% are anchored onto 27 scaffolds, and recovers 98.5% of Hymenoptera-specific single-copy orthologs. Our annotation contains 17,426 protein-coding genes, with a BUSCO completeness of 97.4%.

Previously published single insect genomes have used either Nanopore or PacBio sequencing, sometimes coupled with whole genome amplification or DNA extraction tailored to small starting material. We used standard extraction protocol from haploid tissue with PacBio sequencing and produced haploid reference genome reaching similar BUSCO and N50 statistics as previous single insect genomes (supplementary table 4).

This work provides a crucial resource to study speciation and contemporary hybridization in the *Formica rufa* group, as well as the evolution of extreme sociality. It will also enable new approaches on the genomics of hybridization in this fascinating system. Finally, it also demonstrates that high-quality arthropod genomes can be assembled from single individuals using standard, cost-effective protocols.

## ACKNOWLEDGMENTS

We thank Alan Brelsford for sharing the *F. selysi* assembly and we acknowledge Daniel Blande, Dominik Laetsch, Alex Mackintosh and Yannick Wurm for feedback. We are grateful to CSC – IT Center for Science, Finland, for computational resources. This work was performed under the umbrella of the Global Ant Genomic Alliance and was supported by an HiLIFE fellowship and an Academy of Finland grant no. 309580 to JK.

## AUTHOR CONTRIBUTIONS

PN & JK designed the project. PN, JB & JK performed sampling. JB & JK generated the RNAseq data. PN carried all analyses and drafted the manuscript, which all authors revised for important intellectual content.

## DATA ACCESSIBILITY

The genome assembly has been deposited at the European Nucleotide Archive under the study PRJEB41943. The gene annotation, associated protein sequences and RNAseq data used for annotation purposes are available on Figshare (doi: 10.6084/m9.figshare.c.5332442.v1 & 10.6084/m9.figshare.c.5277767).

