## Supplementary material for "Cost-effective long-read assembly of a hybrid *Formica aquilonia* × *Formica polyctena* wood ant genome from a single haploid individual"


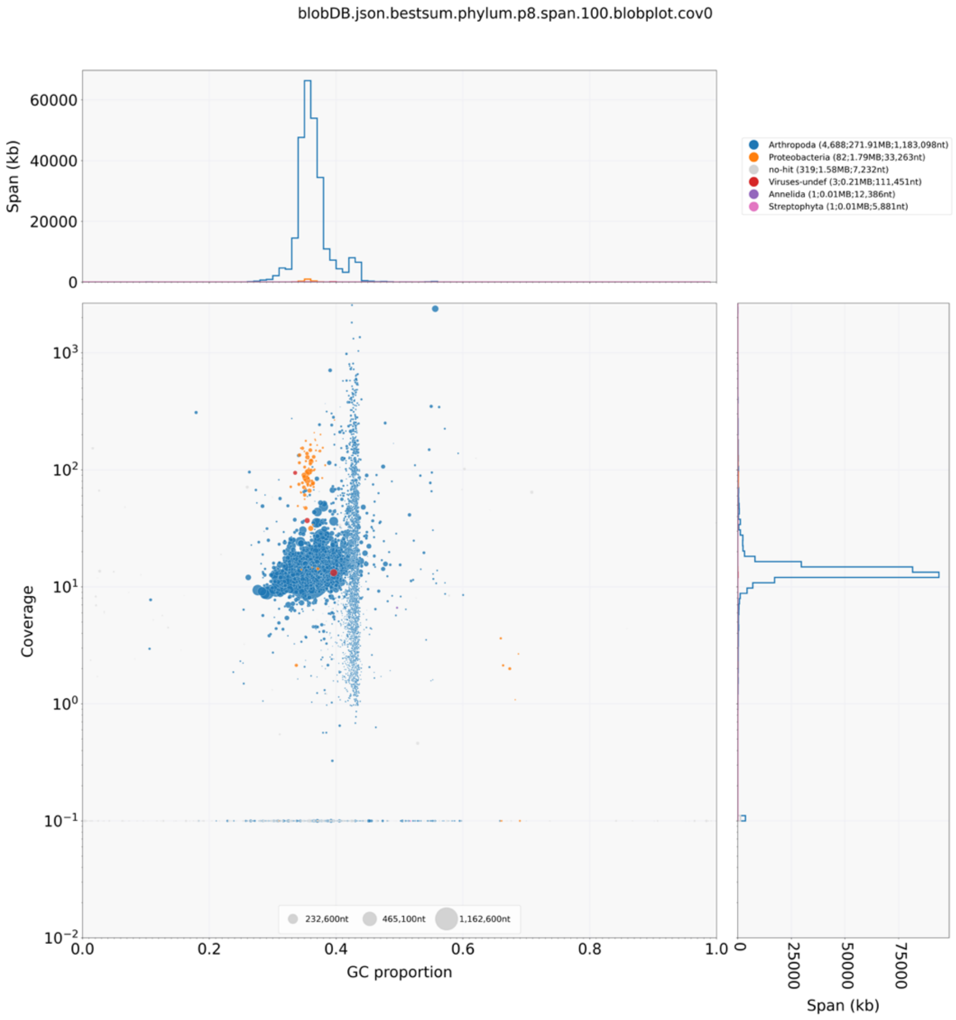


**Supplementary figure S1.** Results of the contamination assessment using Blobtools and the PacBio data.

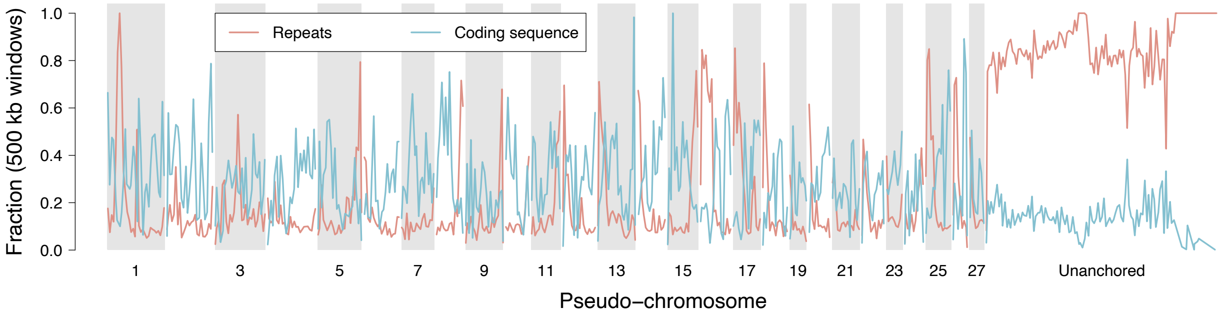


**Supplementary figure S2.** Fraction of repeats and coding sequences computed genome-wide in non-overlapping 500kb windows. Note that the unanchored portion of the genome is repeat-rich and gene-poor.

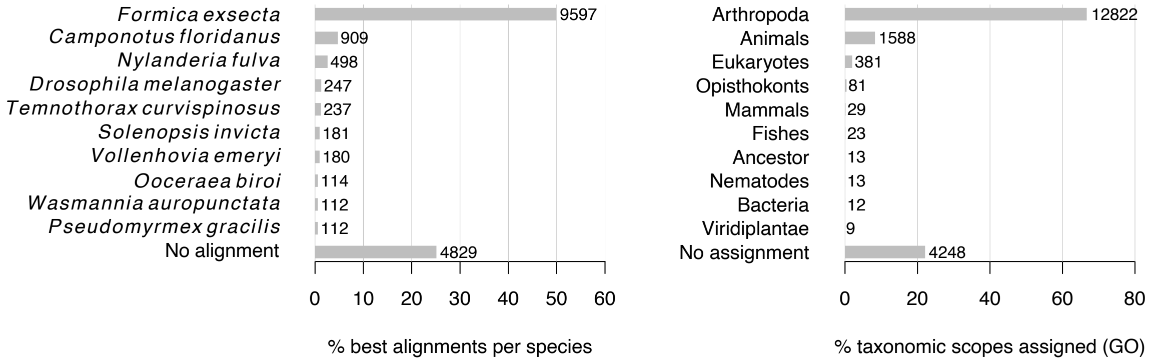


**Supplementary figure S3.** Results of the functional annotation pipeline. The left panel shows the first ten species with the highest number of best hits after similarity search using Diamond and both RefSeq nr and Uniprot databases. Apart from the fruit fly *D. melanogaster* (ranked 4th), all ten most abundant species are ants. Overall, 63.4% of the proteome has best hits within one of these species (25.1% did not get any proper alignment). On the right panel are displayed the number and fraction of transcripts assigned to the first ten Gene Ontology taxonomic scopes using EggNOG.

**Supplementary table S1.** Raw assembly statistics. All sizes are given in base pairs, and statistics are computed assuming a haploid genome size of 323 Mb (see main text).

| Assembler | Subread size cutoff | Total assembly size | Number of contigs | Longest contig | N50 | L50 | N75 | L75 | NG50 | LG50 | NG75 | LG75 | BUSCO v4.0.5 genome score |
|---|---|---|---|---|---|---|---|---|---|---|---|---|---|
| canu | 0 (none) | 338082226 | 3633 | 3815783 | 260777 | 304 | 93570 | 832 | 260777 | 304 | 93947 | 831 | C:97.5%[S:91.8%,D:5.7%],F:1.0%,M:1.5%,n:5991 |
| wtdbg2 | 0 (none) | 349682633 | 11615 | 781157 | 59862 | 1255 | 23343 | 3721 | 70590 | 1051 | 28457 | 2946 | C:79.5%[S:79.1%,D:0.4%],F:4.6%,M:15.9%,n:5991 |
| wtdbg2 | 10000 | 280368309 | 5098 | 4663969 | 1125534 | 68 | 191235 | 231 | 689459 | 93 | 57231 | 534 | C:97.1%[S:96.6%,D:0.5%],F:1.3%,M:1.6%,n:5991 |

**Supplementary table S2.** Annotation statistics for alternative Braker runs. The etp run was considered the best based on the BUSCO score, and its output was later filtered to get rid of spurious gene models.

| Braker2 mode | Data | | Total number of gene models | BUSCO v4.0.5 genome score |
|---|---|---|---|---|
| | RNAseq | Protein | | |
| ep | no | yes | 32,500 | C:95.6%[S:94.9%,D:0.7%],F:2.3%,M:2.1%,n:5991 |
| et | yes | no | 33,077 | C:95.5%[S:94.8%,D:0.7%],F:2.0%,M:2.5%,n:5991 |
| etp | yes | yes | 30,068 | C:97.4%[S:96.8%,D:0.6%],F:1.4%,M:1.2%,n:5991 |

**Supplementary table S3.** Assembly and annotation metrics for the 24 ant genomes for which annotation is available on NCBI.

| Organism Scientific Name | Taxonomy id | Assembly Name | Assembly Accession | Source | Contig N50 (bp) | Size (bp) | Submission Date | BUSCO v4.0.5 genome score | Annotation | Gene Count |
|---|---|---|---|---|---|---|---|---|---|---|
| *Lasius niger* | 67767 | ASM104565v1 | GCA_001045655.1 | GenBank | 17048 | 236236391 | 2.7.2015 | C:89.2%[S:88.5%,D:0.7%], F:7.9%,M:2.9%,n:5991 | INSDC submitter | 18247 |
| *Temnothorax longispinosus* | 300112 | tlon_1.0 | GCA_004794745.1 | GenBank | 30134 | 260681563 | 15.4.2019 | C:95.5%[S:95.0%,D:0.5%], F:2.4%,M:2.1%,n:5991 | INSDC submitter | 13028 |
| *Atta cephalotes* | 12957 | Attacep1.0 | GCF_000143395.1 | RefSeq | 14798 | 317671980 | 6.7.2010 | C:94.3%[S:94.3%,D:0.0%], F:2.8%,M:2.9%,n:5991 | NCBI Annotation Release 100 | 11011 |
| *Pogonomyrmex barbatus* | 144034 | Pbar_UMD_V03 | GCF_000187915.1 | RefSeq | 11605 | 235645958 | 4.2.2011 | C:95.4%[S:95.3%,D:0.1%], F:2.6%,M:2.0%,n:5991 | NCBI Annotation Release 101 | 12761 |
| *Solenopsis invicta* | 13686 | Si_gnH | GCF_000188075.2 | RefSeq | 21162 | 398979080 | 1.8.2018 | C:97.2%[S:96.9%,D:0.3%], F:1.4%,M:1.4%,n:5991 | NCBI Annotation Release 103 | 16814 |
| *Acromyrmex echinatior* | 103372 | Aech_3.9 | GCF_000204515.1 | RefSeq | 80630 | 295944863 | 3.5.2011 | C:98.2%[S:97.9%,D:0.3%], F:0.8%,M:1.0%,n:5991 | NCBI Annotation Release 100 | 12253 |
| *Linepithema humile* | 83485 | Lhum_UMD_V04 | GCF_000217595.1 | RefSeq | 35858 | 219500750 | 10.6.2011 | C:98.2%[S:98.0%,D:0.2%], F:0.8%,M:1.0%,n:5991 | NCBI Annotation Release 100 | 12952 |
| *Vollenhovia emeryi* | 411798 | V.emery_V1.0 | GCF_000949405.1 | RefSeq | 32417 | 287900827 | 6.3.2015 | C:98.2%[S:97.8%,D:0.4%], F:1.0%,M:0.8%,n:5991 | NCBI Annotation Release 100 | 15674 |
| *Wasmannia auropunctata* | 64793 | wasmannia.A_1.0 | GCF_000956235.1 | RefSeq | 37912 | 324120201 | 17.3.2015 | C:97.7%[S:97.4%,D:0.3%], F:1.2%,M:1.1%,n:5991 | NCBI Annotation Release 100 | 15321 |
| *Dinoponera quadriceps* | 609295 | ASM131382v1 | GCF_001313825.1 | RefSeq | 29911 | 259665865 | 13.10.2015 | C:97.7%[S:97.5%,D:0.2%], F:1.1%,M:1.2%,n:5991 | NCBI Annotation Release 100 | 11907 |
| *Atta colombica* | 520822 | Acol1.0 | GCF_001594045.1 | RefSeq | 15290 | 291257934 | 25.3.2016 | C:98.4%[S:98.2%,D:0.2%], F:0.8%,M:0.8%,n:5991 | NCBI Annotation Release 100 | 11174 |
| *Trachymyrmex zeteki* | 64791 | Tzet1.0 | GCF_001594055.1 | RefSeq | 52131 | 267973152 | 31.3.2016 | C:98.3%[S:98.2%,D:0.1%], F:0.8%,M:0.9%,n:5991 | NCBI Annotation Release 100 | 12066 |
| *Cyphomyrmex costatus* | 456900 | Ccosl1.0 | GCF_001594065.1 | RefSeq | 74312 | 300316566 | 25.3.2016 | C:98.3%[S:97.7%,D:0.6%], F:0.7%,M:1.0%,n:5991 | NCBI Annotation Release 100 | 12460 |
| *Trachymyrmex cornetzi* | 471704 | Tcor1.0 | GCF_001594075.1 | RefSeq | 29356 | 369438293 | 25.3.2016 | C:97.9%[S:97.6%,D:0.3%], F:1.1%,M:1.0%,n:5991 | NCBI Annotation Release 100 | 13851 |
| *Trachymyrmex septentrionalis* | 34720 | Tsep1.0 | GCF_001594115.1 | RefSeq | 14962 | 291747019 | 25.3.2016 | C:98.3%[S:98.1%,D:0.2%], F:0.7%,M:1.0%,n:5991 | NCBI Annotation Release 100 | 12049 |
| *Pseudomyrmex gracilis* | 219809 | ASM200609v1 | GCF_002006095.1 | RefSeq | 38830 | 282776121 | 23.2.2017 | C:98.3%[S:97.8%,D:0.5%], F:0.7%,M:1.0%,n:5991 | NCBI Annotation Release 100 | 12655 |
| *Temnothorax curvispinosus* | 300111 | ASM307098v1 | GCF_003070985.1 | RefSeq | 38942 | 303539295 | 24.4.2018 | C:93.9%[S:88.1%,D:5.8%], F:2.4%,M:3.7%,n:5991 | NCBI Annotation Release 100 | 17453 |
| *Harpegnathos saltator* | 610380 | Hsal_v8.5 | GCF_003227715.1 | RefSeq | 911506 | 334536844 | 14.6.2018 | C:98.4%[S:97.6%,D:0.8%], F:0.7%,M:0.9%,n:5991 | NCBI Annotation Release 102 | 14340 |
| *Camponotus floridanus* | 104421 | Cflo_v7.5 | GCF_003227725.1 | RefSeq | 1278439 | 284009182 | 14.6.2018 | C:98.9%[S:98.4%,D:0.5%], F:0.4%,M:0.7%,n:5991 | NCBI Annotation Release 102 | 14020 |
| *Formica exsecta* | 72781 | ASM365146v1 | GCF_003651465.1 | RefSeq | 24299 | 277633851 | 15.10.2018 | C:97.3%[S:95.0%,D:2.3%], F:1.5%,M:1.2%,n:5991 | NCBI Annotation Release 100 | 13725 |
| *Ooceraea biroi* | 2015173 | Obir_v5.4 | GCF_003672135.1 | RefSeq | 3735272 | 223876465 | 23.10.2018 | C:98.0%[S:97.5%,D:0.5%], F:0.7%,M:1.3%,n:5991 | NCBI Annotation Release 100 | 14128 |
| *Nylanderia fulva* | 613905 | TAMU_Nfulva_1.0 | GCF_005281655.1 | RefSeq | 320712 | 375107333 | 13.5.2019 | C:96.1%[S:94.2%,D:1.9%], F:1.4%,M:2.5%,n:5991 | NCBI Annotation Release 100 | 18917 |
| *Odontomachus brunneus* | 486640 | Obru_v1 | GCF_010583005.1 | RefSeq | 22002 | 393036571 | 13.2.2020 | C:95.7%[S:95.1%,D:0.6%], F:2.1%,M:2.2%,n:5991 | NCBI Annotation Release 100 | 13965 |
| *Monomorium pharaonis* | 307658 | ASM1337386v2 | GCF_013373865.1 | RefSeq | 1861574 | 325506644 | 14.8.2020 | C:98.1%[S:95.9%,D:2.2%], F:0.9%,M:1.0%,n:5991 | NCBI Annotation Release 102 | 17083 |

**Supplementary table S4.** Comparison of some recent single-individual-based arthropod genome assemblies.

| Study | Species | Extraction | Input amount | Sequencing Platform | Contig number | Assembly size (Mb) | N50 (Mb) | Scaffolding | Complete BUSCOs |
|---|---|---|---|---|---|---|---|---|---|
| Kingan, Heaton et al. 2020 | mosquito *Anopheles coluzzii* | Modified Qiagen MagAttract protocol | 100 ng | PacBio | 206 | 251 | 3.47 | No | 98.00 % |
| Kingan, Urban et al. 2020 | lanternfly *Lycorma delicatula* | Modified Chromium™ Genome Protocol | 5 µg | PacBio | 2927 | 2 252 | 1.52 | No | 96.80 % |
| Adams et al. 2020 | fruit fly *Drosophila melanogaster* | Qiagen MagAttract | 78.3 ng | Illumina, Nanopore, HiC | NA | 111 | 26.3 | Hi-C | 95.20 % |
| Ye et al. 2020 | parasitoid wasp *Habrobracon hebetor* | TIANamp Micro DNA Kit with WGA | 20 ng | Nanopore | 765 | 132 | 1.63 | No | 99.00 % |
| This study | wood ant *Formica aquilonia* × *F. polyctena* hybrid | standard SDS | 9.89 µg | PacBio | 4687 | 272 | 1.16 | Reference-guided | 98.50 % |
